# Amyloid β-peptide impacts on glucose regulation are dependent on apolipoprotein E genotype

**DOI:** 10.1101/2022.06.22.497222

**Authors:** Jin Hee Sung, Yang Ou, Steven W. Barger

## Abstract

The apolipoprotein E gene (*APOE*) constitutes the greatest genetic risk factor for Alzheimer’s disease, wherein the ε4 allele confers a dramatically elevated risk compared to the more common ε3 allele. Biological mechanisms that differ across these alleles have been explored in mouse models wherein the murine *Apoe* gene has undergone targeted replacement with sequences encoding human ApoE3 or -4 (ApoE-TR mice). Results with such models have indicated that the two variants of ApoE produce differential effects on energy metabolism, including metabolic syndrome. However, glucose regulation has not been compared in ApoE-TR mice with and without Aβ accumulation. We crossed ApoE3- and ApoE4-TR mice with a transgenic line that accumulates human Aβ_1-42_. In male ApoE3-TR mice, introduction of Aβ caused aberrations in glucose tolerance and membrane translocation of astrocytic glucose transporter 1. Phosphorylation of Tau at AD-relevant sites was correlated with glucose intolerance. These effects appeared independent of insulin dysregulation and were not observed in females. In ApoE4-TR mice, the addition of Aβ had no significant effects due to a trend toward perturbation of the baselines. Thus, metabolic changes may have a larger interaction with AD pathology and its consequences in individuals who do not carry an *APOE* ε4 allele. The fact that ApoE4 generally failed to exacerbate the effects of Aβ on glucose further highlights the growing distinction between the glycemic effects of Aβ versus those of peripheral insulin resistance.

## Introduction

Alzheimer’s disease (AD) is a chronic and irreversible brain disorder that involves memory loss and decline of cognitive ability. Risk for developing AD is elevated in a dose-dependent manner by the ε4 allele of the gene (*APOE*) for apolipoprotein E (ApoE), relative to the more common ε3 allele. A wealth of genetic, biochemical, and neuropathological evidence indicates that CNS accumulation of amyloid β-peptide (Aβ)—either as plaques or in a more diffuse, even soluble, form—is the precipitating etiological event for AD. Accumulation of Aβ is higher in individuals with an *APOE* ε4 allele. This relationship has led to the hypothesis that *APOE* ε4 is associated with higher rates of AD because its protein product (ApoE4) promotes Aβ aggregation or retention in the brain. However, other mechanisms are possible, and considerable evidence indicates that the protein product of the *APOE* ε4 allele (ApoE4) enters the cell nucleus and binds specific gene enhancer elements (Lima et al., 2020; Parcon et al., 2018).

One of the earliest events in AD pathogenesis after CNS accumulation of Aβ is a decline in cerebral metabolism of glucose (CMR_glc_) (Jack et al., 2000). The decline in CMR_glc_ is highly correlated with cognitive impairment, and it has impressive specificity as a predictor of progression from mild cognitive impairment (MCI) to frank dementia (Shen et al., 2019; Smailagic et al., 2018) or from normal cognition to MCI (de Leon et al., 2001; Mosconi et al., 2008; Reiman et al., 1998). We recently determined that CMR_glc_ contributes to somatic glucose tolerance, as a transgenic mouse line that accumulates Aβ in the CNS has both reduced CMR_glc_ and impaired glucose tolerance. This appears to result from a decrease in the fraction of glucose transporter 1 (GLUT1) subcellularly localized to the plasma membrane of astrocytes (or other CNS parenchymal cells) (Hendrix et al., 2020). This effect on the 45-kDa form of GLUT1 was also observed in human cerebrum and was the only significant alteration in glucose transporters we identified; reduction of GLUT3, the only CNS transporter that is unique to neurons, does not suppress CMR_glc_ (Stuart et al., 2011).

Persons homozygous for the ε4 allele of *APOE* show an age-related drop in CMR_glc_ in the absence of—or, implicitly, prior to—cognitive impairment (Reiman et al., 1996; Reiman et al., 1998; Small et al., 1995). Female mice carrying targeted replacement (TR) of the murine *Apoe* gene with sequences encoding human ApoE4 show impaired glucose tolerance compared to ApoE3-TR mice (Koren-Iton et al., 2020), but this difference may disappear when compounded by the pathologies associated with diet-induced obesity or Aβ accumulation (Christensen and Pike, 2019; Koren-Iton et al., 2020). The latter manipulation was performed through transgenesis for the full amyloid precursor protein (APP), which has marked effects on peripheral glucose regulation (Botteri et al., 2018; Kulas et al., 2018; Needham et al., 2008; Tu et al., 2012). We sought to determine the impact of human ApoE4 on glucose regulation in a mouse model that suffers CNS Aβ accumulation without an APP transgene. Unlike prior studies, we directly compared ApoE-TR mice with and without human Aβ expression. ApoE3-TR mice responded to the CNS accumulation of Aβ with hyperphosphorylation of Tau that correlated with impairments in glucose tolerance and trafficking of the 45-kDa GLUT1. These effects were not observed in ApoE4-TR mice due to pathological trends in the absence of the Aβ transgene.

## Materials and Methods

### ApoE-targeted replacement mice

Mice homozygous for ApoE3- or ApoE4-TR loci (Taconic Biosciences) were assessed alongside ApoE-TR mice from the same breeding colony that were also carrying an Aβ_1-42_ transgene (Aβ-Tg) hemizygously, the latter generated by crossing the ApoE-TR lines with the BRI-Aβ42 line (McGowan et al., 2005). Mice were maintained on a 12-h light/dark cycle at 23 °C with free access to food and water. All animal experiments were conducted according to the Institutional Animal Care and Use Committee of the Central Arkansas Veterans Healthcare System (CAVHS), which is certified by the Association for Assessment and Accreditation of Laboratory Animal Care International (AAALAC). Mice were anesthetized in CO_2_ chamber. Left brains were collected and stored at -80 for biochemical assays.

### Genotyping

Blood samples from tail nicks were collected on Whatman FTA card (Millipore Sigma, St. Louis, MO, USA). Total genomic DNA from blood was isolated using REDExtract-N-Amp Blood PCR Kit (Millipore Sigma) following the manufacturer’s instructions. PCR was performed on the extracted DNA with the following primers: BRI-Aβ42 Forward: 5’-AAG GCT GGA ACC TAT TTG CC-3′, Reverse: 5’-TAG TGG ATC CCT ACG CTA TG-3′; APOE-TR Forward: 5’-GGT AGC TAG CCT GGA CGG-3′, Reverse: 5’-GCT AGA ACC AGC AGA GAC CC-3′; *Apoe*-WT Forward: 5’-GTC TCG GCT CTG AAC TAC TAT-3′, Reverse: 5’-GCA AGA GGT GAT GGT ACT CG-3′.

### GTT and ITT

Glucose tolerance test (GTT) was performed at 7 weeks or 12 months of age. Mice fasted for 6 h, and then D-(+)-glucose (2 g/kg) was injected intraperitoneally. Concentration of glucose in tail-vein blood ([Glc]_b_] was assayed before injection and at 15, 30, 60 and 120 min after injection using an AlphaTRAK® Glucose Meter (Zoetis Inc, Kalamazzo MI). Area under the curve was calculated as total AUC (tAUC), which compares absolute values of [Glc]_b_. Insulin tolerance test (ITT) performed to assess insulin sensitivity at 7 weeks of age. Non-fasted mice were transferred to individual cages injected with 1 U/kg insulin (Humulin R, Lily) intraperitoneally. Blood glucose was measured by the same method as for GTT. Because ITT produces a downward deflection from baseline, integration of these values was performed as area over the curve (AOC), measured from the initial [Glc]_b_.

### Western blot analysis and Wes immunoassay

Harvested brain tissue was stored at -80 °C and then chilled further in liquid N_2_ prior to being pulverized with a liquid N_2_-chilled mortar and pestle. For conventional western blot analysis, pulverized tissue was combined with RIPA buffer (50 mM Tris-HCl, 150 mM NaCl, 1% NP-40, 0.5% sodium deoxycholate, 0.1% SDS, 1 mM Na_3_VO_4_, 1 mM NaF) containing 5 mg/mL protease inhibitors (Pierce Mini Tablets, EDTA-free), pH 7.5) and sonicated on ice. The lysates were quantified for protein with the Pierce BCA protein assay kit (Thermo Fisher Scientific). Total protein (30 μg) per lane was separated by SDS-PAGE using 4–15% Tris-glycine gels (Bio-Rad Laboratories) and transferred to PVDF membrane (Millipore Sigma). The blots were blocked with SuperBlock Blocking Buffer (Thermo Fisher Scientific) for 1 h at room temperature and then incubated with primary antibody overnight at 4 °C: AT8, detecting phosphorylated Tau (Thermo Fisher Scientific, MN1020, 1:250), a phospho-independent anti-Tau antibody (Cell Signaling, #46687, 1:1000), or an anti-IL1β antibody (ABclonal, #A16228, 1:1000). The membranes were washed in Tris-buffered saline containing 0.1% Tween-20 (TBST) and then exposed to secondary antibody: anti-rabbit and anti-mouse HRP-conjugated (Thermo Fisher Scientific). After washing in TBST, the membranes were reacted with enhanced chemiluminescent substrate (Thermo Fisher Scientific). Protein bands were visualized by exposure to x-ray film and quantification performed with ImageJ.

For analysis of GLUT1, membrane fractionation was performed with a Minute™ Plasma Membrane Protein Isolation and Cell Fractionation Kit (Invent Biotechnologies, Plymouth MN), which allows separation of tissue homogenates into plasma membrane (PM), cytosolic, and organellar fractions. As previously described (Hendrix et al., 2020), these fractions were analyzed on a SimpleWes™ capillary electrophoresis device (Protein Simple), which provides requisite separation of the molecular-weight moieties and quantitation. The value obtained for the 45-kDa GLUT1 in each fraction was multiplied by the dilution factor of each, then the PM value was divided by the total of all fractions.

### ELISA

After a 6-h fast, tail-vein blood was collected at 0 and 30 min relative to an i.p. injection of 2 g/kg glucose (as per GTT). Blood samples were allowed to coagulate and then centrifuged at 17,000 *g* for 15 min at 4 °C. Serum samples were stored at -80 °C. Insulin ELISA was performed using an Ultra-Sensitive Mouse Insulin ELISA Kit (Crystal Chem) according to manufacturer’s instructions.

For cerebral Aβ ELISA, pulverized brain samples were weighed before extraction. Brain tissue were homogenized in 15 times ice-cold extraction buffer [Tris-buffered saline (TBS) containing 1 mM Na_3_VO_4_, 1 mM NaF, and 5 mg/mL protease inhibitors (Pierce Mini Tablets, EDTA-free)] and centrifuged at 100,000 *g* for 60 min at 4 °C. The supernatants were stored for determination of water-soluble Aβ. The pellets were washed in TBS and suspended in 50 μl extraction buffer with 1% Triton X-100. The suspensions were mixed using a gentle rotator for 30 min 4 °C and centrifuged at 100,000 *g* for 60 min at 4 °C. These supernatants were stored for determination of detergent-soluble Aβ. The pellets were washed in TBS and resuspended in extraction buffer containing 70% formic acid. The samples were mixed on a rotator for 2 h at room temperature. After mixing, the suspensions were centrifuged at 100,000 *g* for 60 min at 4 °C. These supernatants were stored for acid-soluble Aβ analysis. All supernatants were stored at -80 °C prior to ELISA. Aβ ELISA was performed using a human amyloid β (aa1-42) immunoassay Quantikine ELISA kit (R&D Systems, Inc.).

### Primary culture of astrocytes

Astrocytes were isolated from postnatal-day 1–3 Sprague-Dawley rat pups or mouse pups of wild-type, ApoE3-TR, and ApoE4-TR genotypes. Pups anesthetized at by hypothermia, sanitized with 70% ethanol, and transferred into HEPES-buffered Hank’s balanced salt solution (HBSS) (Millipore Sigma). Brains were removed and the cerebral hemispheres removed from midbrain and brainstem. Meninges and olfactory bulbs were removed. Cerebral hemispheres were diced with a scalpel blade and incubated in 0.1 mg/ml trypsin (Thermo Fisher Scientific) in HBSS for 8 min at room temperature. The tissues were rinsed with HBSS and exposed to 1.5 mg/ml trypsin inhibitor (Thermo Fisher Scientific) for 10 min at room temperature. After rinsing the tissue with HBSS, the tissue was triturated and plated in tissue culture flasks with minimal essential medium (Earle’s salts; MEM) containing 10% fetal bovine serum (FBS). After 10-14 days, the loosely attached microglia were dislodged by striking the flask abruptly and washing with phosphate-buffered saline (PBS). The remaining astrocytes were then trypsinized for subculturing. For subcellular fractionation, cells were plated in 150-mm plates (3 per data point). Once confluent, the culture medium was replaced with Dulbecco’s modified Eagle’s medium (DMEM) containing 0.5% FBS with 5 or 0.2 mM glucose for 24 h.

Human astrocytes were cultured from superior temporal gyrus of de-identified autopsy waste material that was discarded by the Department of Pathology at UAMS. The material was obtained within 6 h of death and was processed as described above for rodent cerebral tissue with the exception that trypsinization was performed for 40 min in 0.02 mg/ml trypsin. This use of autopsy waste was reviewed by the UAMS Institutional Review Board and determined not to constitute human subjects research.

### Glucose uptake assay

Aβ_1-42_ (rPeptide) was prepared by first dissolving in hexafluoroisopropanol (HFIP) at 1 mM for 3-4 h to remove any pre-existing fibrils. The HFIP was removed by evaporation, and the peptide was dissolved in anhydrous dimethyl sulfoxide at 2 mM, then diluted to 150 μM in ice-cold PBS. The solution was incubated at 4 °C for 24 h to allow for the formation of oligomers.

Subcultured primary astrocytes were seeded in 24-well plates and grown to confluency. Once confluent the growth medium was replaced with MEM containing 0.5% FBS with 2.5 μM Aβ_1-42_ or vehicle for 4 h prior to harvest. The medium was then replaced with DMEM lacking glucose, containing 10 μCi/ml [^3^H]2-deoxyglucose and 1 μCi/ml [^14^C]sucrose. After 30 min, the medium was removed and stored for scintillation counting. The cells were washed twice with ice-cold DMEM containing 25 mM glucose then lysed via a freeze-thaw cycle in 10 mM Tris-HCl (pH 7.2) containing 0.2% sodium dodecyl sulfate. One aliquot was subjected to scintillation counting and another to protein determination by BCA reaction. The ^14^C cpm/ml in medium was used to determine the volume of extraneous medium contributing to each cell lysate, and ^3^H cpm corresponding to this volume were subtracted from the total in the lysate. Finally, the remaining ^3^H cpm were normalized to the protein content in the well.

### Experimental design and statistical analysis

Younger male mice (7-9 weeks) numbered as follows: ApoE3-TR: 7, ApoE4-TR 11, ApoE3-TR/Aβ-Tg: 6, ApoE4-TR/Aβ-TR: 7; in some analyses, mice were excluded by a random-number generator to achieve N=6 for the sake of convenient loading of electrophoresis apparatus. For older males attrition limited sample size to 4; 1-year-old females numbered as follows: ApoE3-TR: 5, ApoE4-TR 4, ApoE3-TR/Aβ-Tg: 6, ApoE4-TR/Aβ-TR: 8. Data sets were assessed by one-way ANOVA followed by the *post hoc* test indicated in the figure legends. These calculations were performed with GraphPad Prism 9. Pearson’s correlation coefficient was used to analyze correlations. Values of P less than 0.05 were considered statistically significant.

## Results

### Aβ significantly impairs glucose tolerance in male ApoE3-TR mice

We previously reported that accumulation of Aβ_1-42_ in the CNS of mice is sufficient to impair peripherally measured glucose tolerance without disruption of insulin dynamics. ApoE3- and ApoE4-TR mice were crossed with Aβ-Tg mice, and these were compared to their Aβ-free counterparts regarding glucose and insulin regulation. Male ApoE3-TR/Aβ-Tg mice tested at 12 months of age exhibited impairments in a glucose tolerance test (GTT) compared to ApoE3-TR mice without an Aβ transgene (Fig. 1A). ApoE4-TR mice trended toward an impairment, and this prevented the apparent elevation by Aβ from reaching significance. As we had observed earlier in mice with a wild-type *Apoe* locus (Hendrix et al., 2020), glucose tolerance in females expressing either of the human ApoE variants was unaffected by Aβ accumulation (Fig. 1B). ApoE4-TR females showed a small trend toward impairment compared to ApoE3-TR females, irrespective of Aβ expression, but this did not reach significance. Body weights did not differ between the genotypes in either sex (Table 1). To test the effects of age, additional GTTs were performed on 7-week-old males. Compared to older males, the total area under the curve (tAUC) during the 2-h GTT was 29% lower in the younger ApoE3- and 4TR mice (without the Aβ transgene). Therefore, the results are presented as the percentage of ApoE3-TR values in each respective age group so that relative differences between the genotypes could be more readily compared across the ages (Fig. 1C). The relationships between ApoE variant, Aβ expression, and GTT followed essentially the same pattern in each age group, though Aβ trended toward a greater effect irrespective of ApoE variant. The younger cohort was used for the data presented henceforth.

**Figure 1.**
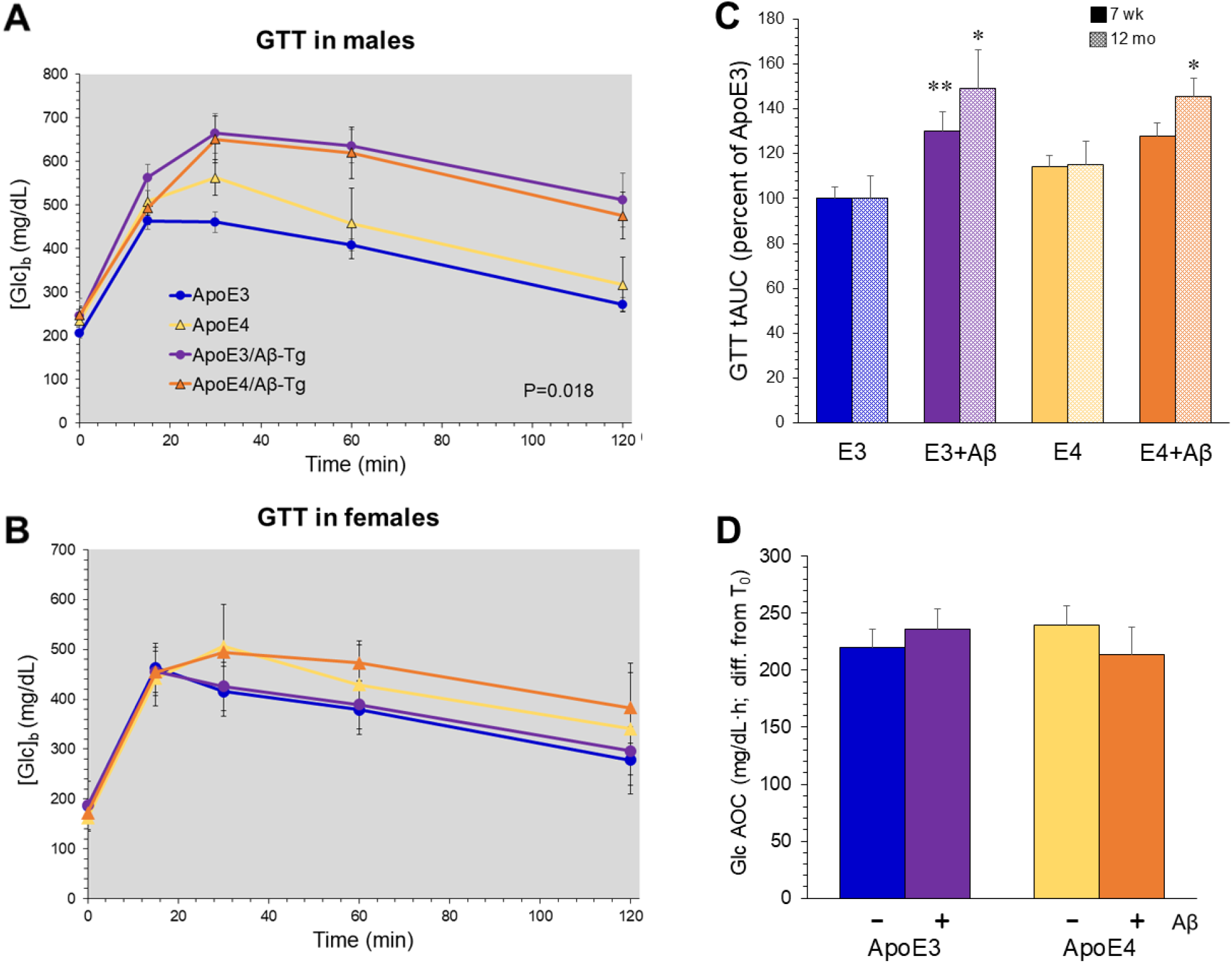
Aβ impairs glucose tolerance in male mice expressing ApoE3. Mice which had undergone targeted replacement (TR) of the endogenous *Apoe* gene with sequences producing human ApoE3 or ApoE4 were crossed with Aβ-Tg mice. They were compared to littermates which did not inherit the Aβ-Tg. GTT was performed on 12-month-old male mice (**A**) or female mice (**B**); p=0.02, ApoE3-TR vs. ApoE3-TR/Aβ-Tg (ANOVA and Bonferroni *post hoc*). **C:** AUC was calculated from GTT values for 7-wk-old and 12-mo-old male mice; results are shown relative to the mean values from ApoE3-TR mice at each age; *p<0.04 vs. ApoE3; **p<0.01 vs. ApoE3 (ANOVA and Tukey *post hoc*). **D:** ITT on males reflected as area over the curve (AOC).

**Table 1.** Body weights.

| Sex | Genotype | Body wt. (g±SEM) |
| --- | --- | --- |
| males | ApoE3 | 43.25 ± 2.070 |
|  | ApoE4 | 46.80 ± 2.881 |
|  | ApoE3/Aβ-Tg | 46.47 ± 0.592 |
|  | ApoE4/Aβ-Tg | 44.12 ± 0.615 |
| females | ApoE3 | 39.06 ± 4.502 |
|  | ApoE4 | 39.32 ± 3.444 |
|  | ApoE3/Aβ-Tg | 34.07 ± 4.823 |
|  | ApoE4/Aβ-Tg | 36.47 ± 3.775 |

As seen in mice with a wild-type *Apoe* locus (Hendrix et al., 2020), the impaired glucose tolerance in ApoE3-TR males was not associated with insulin resistance. ITT showed no differences between any of the genotypes, as reflected in area over the curve in Fig. 1D. Serum levels of insulin were measured after fasting and 30 min after i.p. glucose administration, and no significant differences between the genotypes were detected (Table 2). An effect of Aβ was noted in male ApoE4-TR mice in the form of a basal blood glucose concentration ([Glc]_b_) that remained elevated in the fasted state (Fig. 2 A,B). In the nonfasted state, there was no difference in basal [Glc]_b_ among the groups.

**Table 2.**
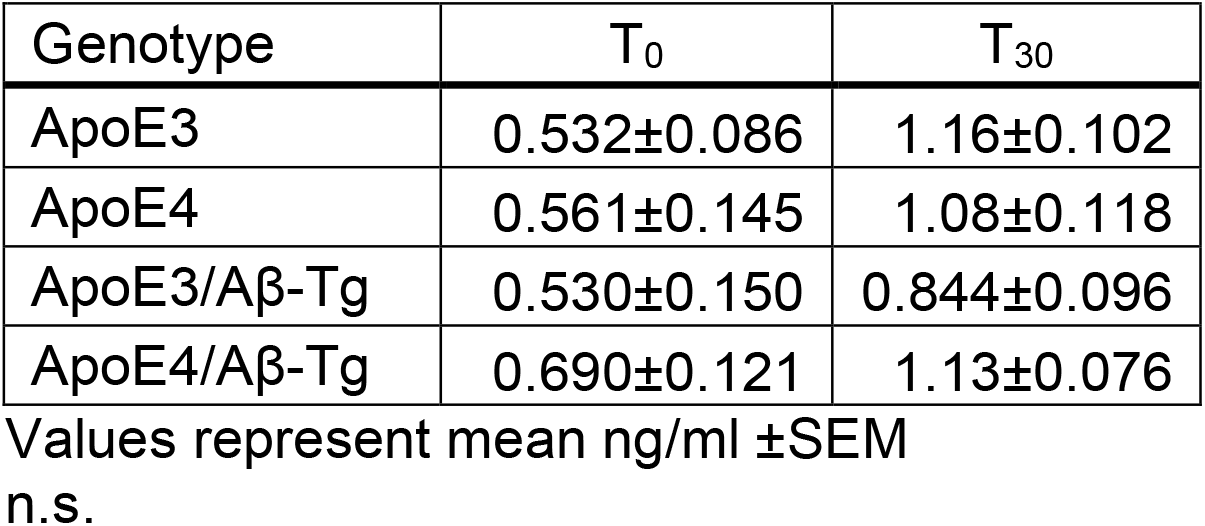
Serum insulin levels.

| Genotype | T<sub>0</sub> | T<sub>30</sub> |
| --- | --- | --- |
| ApoE3 | 0.532±0.086 | 1.16±0.102 |
| ApoE4 | 0.561±0.145 | 1.08±0.118 |
| ApoE3/Aβ-Tg | 0.530±0.150 | 0.844±0.096 |
| ApoE4/Aβ-Tg | 0.690±0.121 | 1.13±0.076 |

Values represent mean ng/ml ±SEM

**Figure 2.**
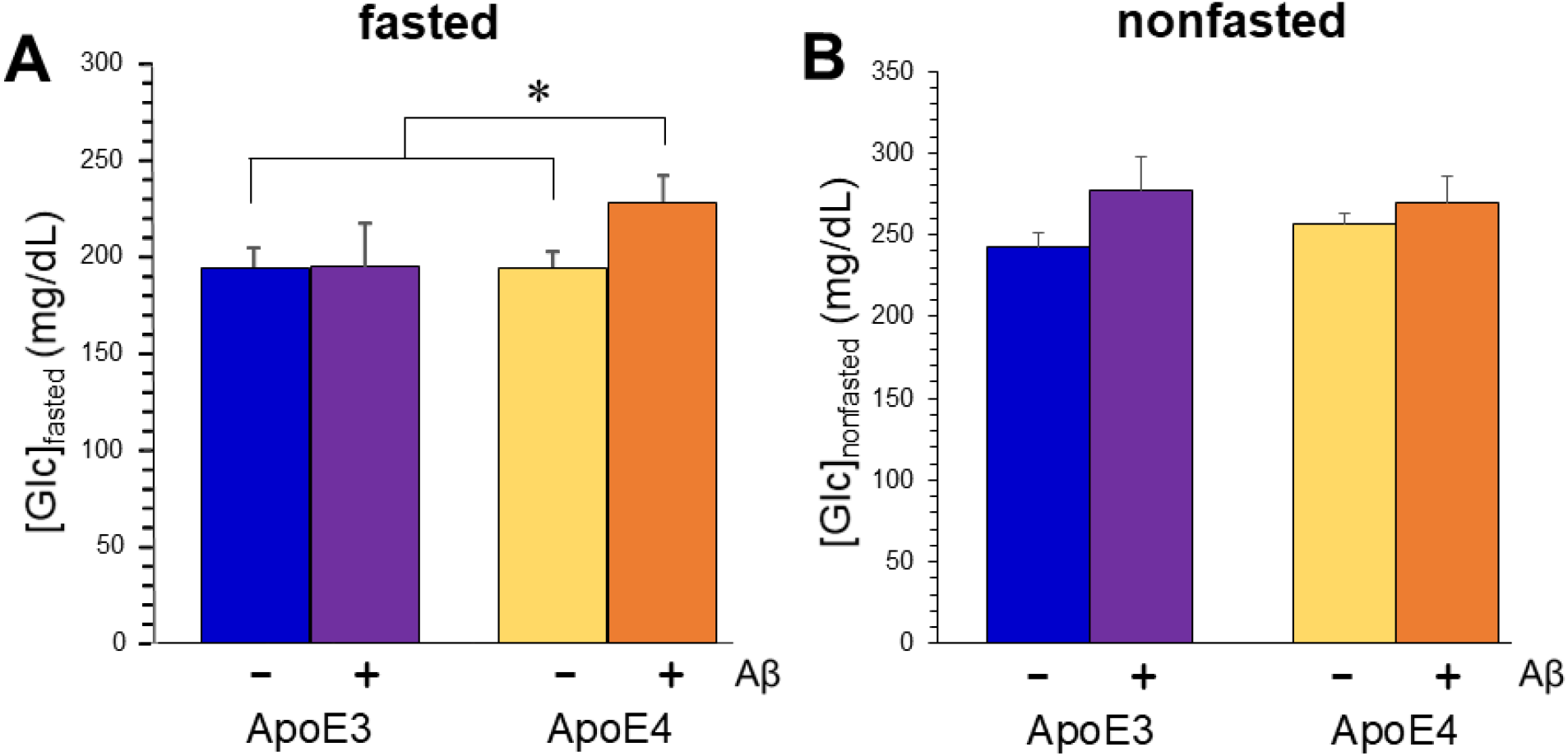
Aβ combines with ApoE4 expression to elevate basal [Glc]_b_. Basal [Glc]_b_ was measured in mice of ApoE3-TR, ApoE3-TR/Aβ, ApoE4-TR, and ApoE4-TR/Aβ genotypes. The mice had either been fasted 5 h (**A**) or not (**B**). Values represent mean ± SEM; *p<0.05 (ANOVA and Tukey *post hoc*).

### Aβ significantly impairs GLUT1 membrane localization in ApoE3-TR mice

The impairments in GTT in male Aβ-Tg mice were previously associated with deficient trafficking of GLUT1 in brain parenchymal cells. Specifically, the proportion of the 45-kDa form of GLUT1 in cortical plasma membrane fractions was lower in these mice, as it was in AD (Hendrix et al., 2020). We examined this property in ApoE-TR males. The results revealed a pattern reciprocal to that of GTT: ApoE3-TR/Aβ-Tg mice exhibited a plasma membrane (PM) fraction of 45-kDa GLUT1 that was significantly lower than that in ApoE3-TR mice; ApoE4-TR mice trended toward a lower value, which obviated a significant difference between these and their ApoE4-TR/Aβ-Tg counterparts (Fig. 3). Total levels of GLUT1 in the four groups did not differ (not shown).

**Figure 3.**
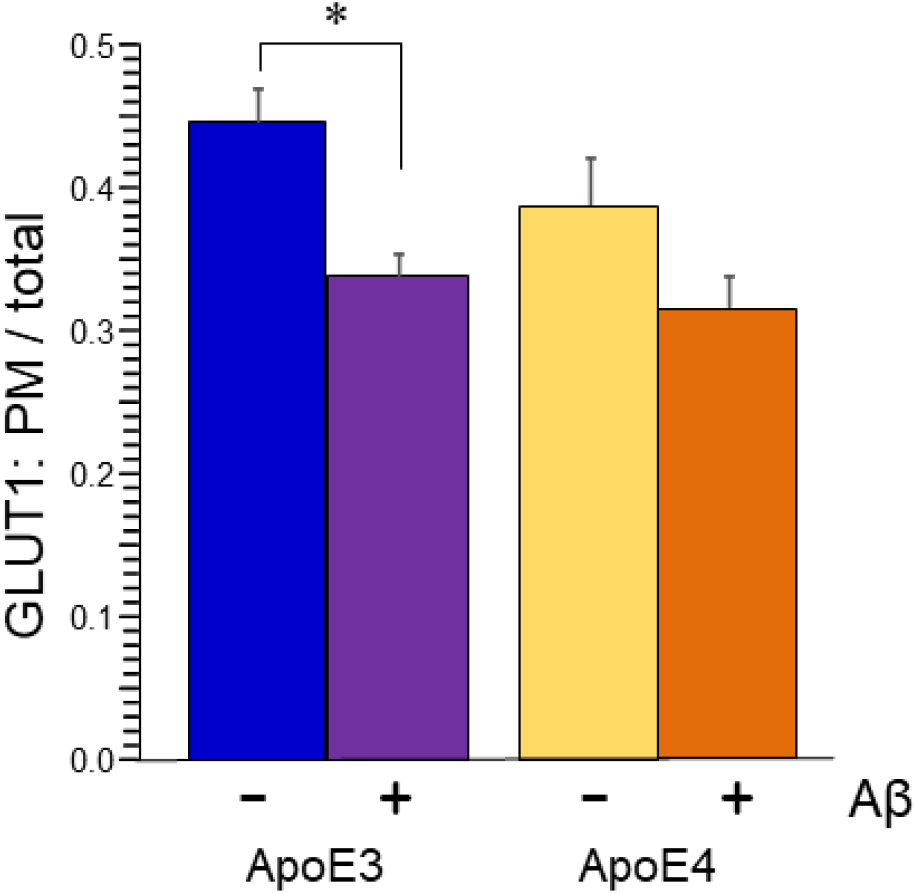
Aβ impairs plasmalemmal localization of GLUT1 in cerebral parenchymal cells of ApoE3-TR mice. Cerebral hemispheres from ApoE3-TR, ApoE3-TR/Aβ, ApoE4-TR, and ApoE4-TR/Aβ genotypes were homogenized and fractionated to isolate plasma membrane. Steady-state levels of the 45-kDa isoform of GLUT1 were determined in this fraction and the remainder of cellular components using SimpleWes™ capillary electrophoresis. Values represent mean ± SEM; *p<0.05 (ANOVA and Tukey *post hoc*).

### Aβ and ApoE4 conspire to alter GLUT1 and glucose uptake in cultured astrocytes

We also assayed the PM fraction of 45-kDa GLUT1 in astrocytes cultured from ApoE3- and ApoE4-TR mice. Basal PM fraction of GLUT1 was not significantly different between ApoE3- and ApoE4-expressing astrocytes under normal culture conditions. As seen in prior studies with wild-type adipocytes and myocytes, reduction of glucose concentration in the culture medium enhanced the amount of GLUT1 in the PM fraction in astrocytes from ApoE3-TR mice (Fig. 4A). In contrast, the PM fraction of GLUT1 was deflected downward, resulting in a significant difference between the genotypes.

**Figure 4.**
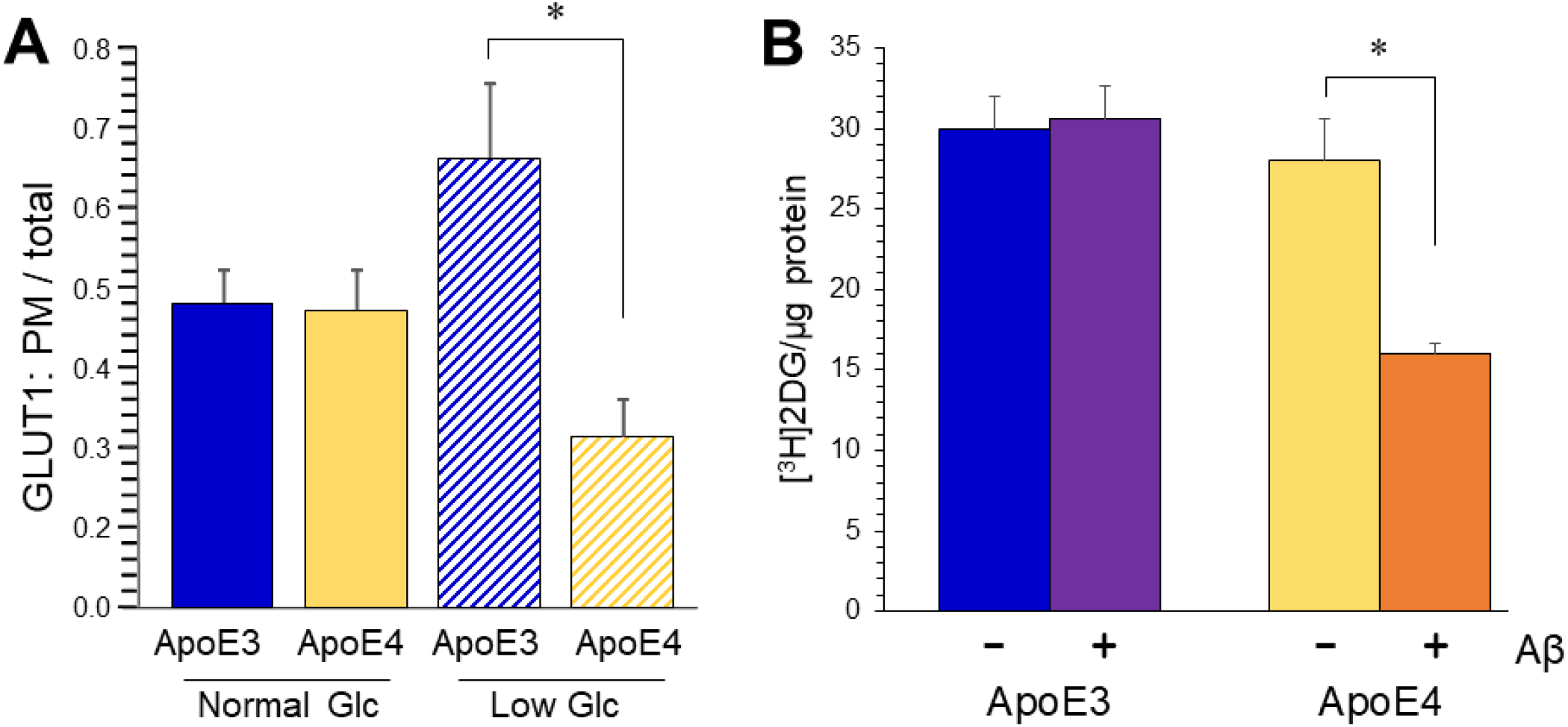
Aβ and ApoE4 conspire to impair glucose transport in cultured astrocytes. Primary astrocyte cultures were prepared from postnatal mice of ApoE3 or ApoE4 mice. **A:** Cells were cultured in DMEM containing 5 mM or 0.2 mM Glc. Homogenates were fractionated to isolate plasma membrane, and levels of the 45-kDa GLUT1 were determined by SimpleWes™. **B:** Cultures were treated with 2.5 μM aggregated Aβ_1-42_. After 20 h, a 30-min uptake of [^3^H]2-deoxyglucose was measured, corrected by inclusion of [^14^C]sucrose and normalized to protein quantity. Values represent mean ± SEM (N=4); *p=0.05 (ANOVA and Tukey *post hoc*).

To test for effects of ApoE variant on glucose uptake, we assayed acute accumulation of [^3^H]2-deoxyglucose (2DG) in cultured astrocytes. Some cultures were exposed to 2.5 μM Aβ_1-42_ for 8 h prior to assay. (This time period was sufficient to down-regulate GLUT1 PM fraction in wild-type astrocytes.) Within a 30-min incubation with [^3^H]2DG, ApoE3- and ApoE4-TR astrocytes had taken up essentially equal amounts compared to ApoE4-TR astrocytes (Fig. 4B). Exposure to Aβ reduced the uptake of 2DG in ApoE4-TR cells, but the rate in ApoE3-TR cells was not significantly impacted by Aβ.

### ApoE4 expression elevates expression of pro-IL-1β without elevating maturation

Activation of inflammatory sequelae is well established in AD, and variant-specific effects of ApoE on these molecules and pathways have been reported (Kloske and Wilcock, 2020; Vitek et al., 2009). Interleukin-1β (IL-1β) is a well-documented component of the neuroinflammatory changes seen in AD and has mechanistic connections to aspects of AD pathology. We measured IL-1β levels in the brains of males with ApoE-TR, with or without the Aβ transgene. Because maturation and release of IL-1β depends on activation of the inflammasome, we prefer to assay this cytokine by western blot rather than ELISA, as the former allows assessment of mobility to distinguish pro-IL-1β (35 kDa) from mature IL-1β (17 kDa). Western-blot analysis of cerebral cortical tissue detected an elevation of pro-IL-1β that was dependent on ApoE4 expression and not Aβ (Fig. 5 A,B). However, mature IL-1β was unaltered by either (Fig. 5 A,C). The latter finding is consistent with a similar recalcitrance of interleukin-1-converting enzyme (ICE; caspase 1) (not shown).

**Figure 5.**
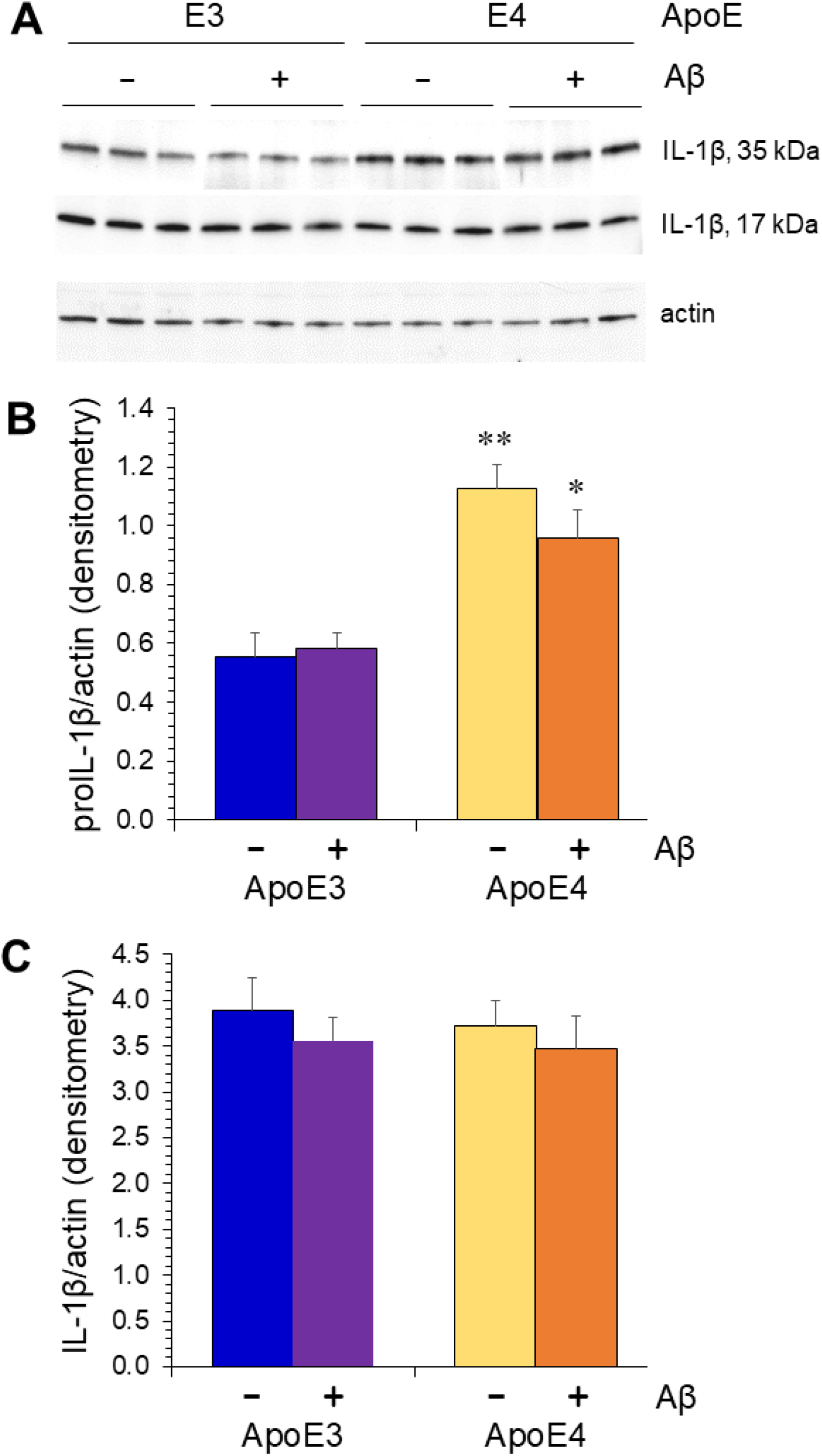
ApoE4 expression is associated with higher levels of pro-IL-1β but not mature IL-1β. Cerebral hemispheres from male mice of ApoE3-TR, ApoE3-TR/Aβ, ApoE4-TR, and ApoE4-TR/Aβ genotypes were homogenized and analyzed by Western blot for IL-1β (**A**); β-actin was quantified as a loading control. The 35-kDa proIL-1β (**B**) and 17-kDa mature IL-1β (**C**) were quantified separately. Values represent mean ± SEM; *p=0.03, **p=0.004 (ANOVA and Tukey *post hoc*).

### Tau hyperphosphorylation mirrors GTT perturbation

To determine if markers related to neurodegeneration were altered by ApoE variant we evaluated the phosphorylation state of Tau in cerebral tissue from ApoE-TR males. Western blots for pathological Tau were performed with the AT8 antibody, which detects Tau phosphorylated at amino acids Ser_202_ and Thr_205_; these sites are phosphorylated in neurofibrillary tangles found in AD and other tauopathies. AT8 immunoreactivity was normalized to results obtained with a phosphorylation-independent Tau antibody. GTT results were tabulated as area under the curve (AUC) to provide an integrated assessment of glucose derangement. Both parameters showed a similar step-wise increase from ApoE3-TR to ApoE4-TR to ApoE3-TR/Aβ-Tg (Fig. 6). Moreover, when the correlation between GTT AUC and AT8/Tau values for each animal, significance was observed for the overall cohort, all ApoE3-TR mice and all mice lacking Aβ (Table 3); analysis of all ApoE4-TR and all Aβ-Tg mice showed no significance.

**Figure 6.**
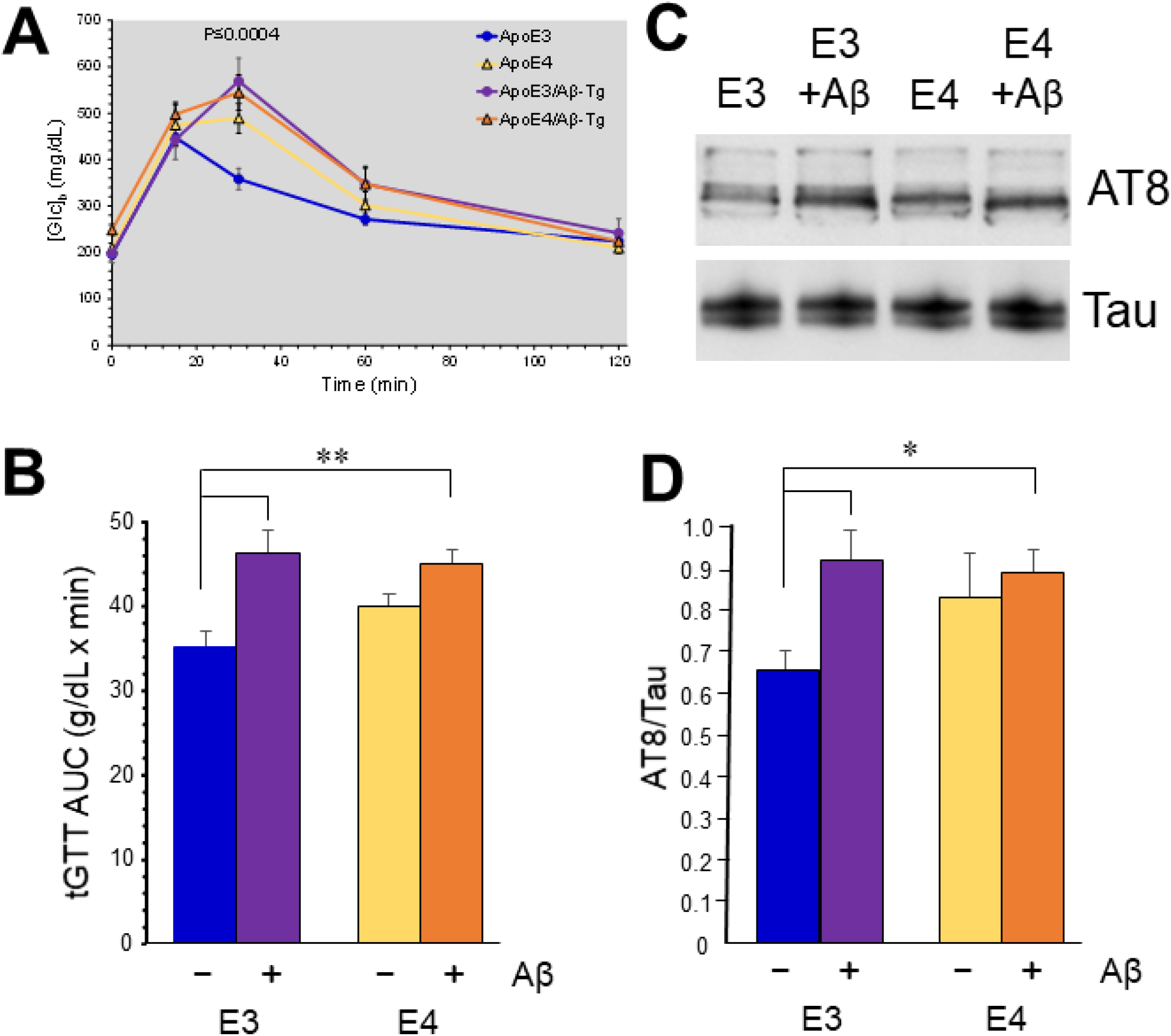
Tau pathology approximates glucose perturbation in Aβ-Tg x ApoE-TR mice. Males of ApoE3-TR and ApoE4-TR with or without the Aβ transgene were compared. **A:** A 2-h glucose tolerance (GTT) assay was performed at 7 weeks of age. **B:** Total area under the curve for the GTT is plotted for each genotype. **C:** Cerebral cortex homogenates were subjected to western-blot analysis with the AT8 antibody or a phospho-independent Tau antibody. **D:** Quantification of AT8/Tau ratio. Values are mean±SEM; *P≤0.05, ** P≤0.01 (ANOVA and Tukey *post hoc*).

**Table 3.** GTT vs. AT8/Tau correlation.

| Group | R | p value |
| --- | --- | --- |
| all mice | 0.46439 | 0.022* |
| all ApoE3 | 0.67955 | 0.021* |
| all ApoE4 | -0.07666 | 0.824 |
| all -Aβ | 0.69847 | 0.013* |
| all +Aβ | -0.28661 | 0.367 |

## Discussion

We previously reported that accumulation of Aβ_1-42_ in the CNS of mice is sufficient to impair glucose tolerance without disrupting the production of or sensitivity to insulin. In fact, as the utilization of glucose in the CNS is largely insulin-independent, impairment of glucose tolerance in the absence of insulin resistance may be a relatively specific marker for reduced CMR_glc_. We evaluated glucose and insulin regulation in mice expressing human ApoE3 and ApoE4 because of the impact of *APOE* genotype on rates of AD and the prior connections of ApoE to energy metabolism. We expected that ApoE4 would exacerbate physiological the effects of Aβ accumulation in the CNS. To effect the latter, we used an Aβ transgene that introduces Aβ_1-42_ into the brain interstitial fluid without an APP transgene. This was deemed an important departure from prior studies due to documented effects of APP on insulin and glucose regulation. Surprisingly, perturbations of glucose tolerance did not show significant cooperative effects of Aβ and ApoE4. Instead, ApoE4 itself tended to shift values in the direction of pathophysiology. In this way, the phenotype observed in ApoE3-expressing mice showed a greater dependence on Aβ. These results are compatible with evidence that ApoE4 impacts glucose utilization independently of AD pathogenesis (e.g., in young adults). If ApoE4 significantly exacerbated Aβ accumulation in this model system, it might be expected that the combination of an Aβ transgene with ApoE4 expression would create additive or perhaps even synergistic effects; but, only basal [Glc]_b_ was affected in this way. Thus, our findings suggest that ApoE4 could have detrimental effects by means other than worsening Aβ accumulation.

A few studies performed in mice combining human ApoE variants and Aβ accumulation have reported glucose dynamics, but none has directly compared ApoE-TR mice with and without the genetic manipulation of Aβ. In addition, prior studies attempted to model AD by overexpressing mutated APP. We and others have noted dramatic effects of APP on insulin and glucose regulation (Botteri et al., 2018; Kulas et al., 2018; Needham et al., 2008; Tu et al., 2012); therefore, we prefer a model of Aβ accumulation that does not depend on overexpression of APP. Pike and colleagues have made considerable study of a model (“EFAD”) wherein ApoE-TR mice are crossed with the 5xFAD line. EFAD mice responded to an obesogenic high-fat, high-sucrose (“western”) diet in a manner reminiscent of the response we observed to Aβ: glucose tolerance was impaired by the diet-induced obesity in ApoE3-expressing mice but not in ApoE4-expressing mice. However, this relationship was shown in females (Christensen and Pike, 2019) but not males (Moser and Pike, 2017), opposite to the sex-dependency of the effects we observed. It is not clear what an increased vulnerability of females means in an animal model that does not experience menopause. While female humans develop AD at a higher rate than males, this is an entirely post-menopausal phenomenon (Riedel et al., 2016; Seshadri et al., 1997).

Several studies have evaluated glucose tolerance in ApoE-TR mice without any expression of human Aβ. Those have generally found no difference between the ApoE variants in GTT assays unless the mice were fed obesogenic diets (Arbones-Mainar et al., 2010; Fleeman et al., 2022; Jones et al., 2019; Mattar et al., 2022). Whereas female C57BL/6 mice with endogenous murine *Apoe* are generally refractory to obesogenic diets (Pettersson et al., 2012), expression of ApoE4 (but not ApoE3) seems to render them vulnerable to glucose intolerance (Jones et al., 2019). A study that exclusively utilized female mice found that, similar to the EFAD results described above, glucose tolerance was impaired by an obesogenic diet in ApoE3-expressing mice but not in ApoE4-expressing mice (Koren-Iton et al., 2020); the converse, with respect to ApoE4 variant, has been reported in males (Arbones-Mainar et al., 2010). One study utilized mass spectrometry to assess plasma ApoE3 and ApoE4 levels in heterozygous human subjects, finding that [Glc]_b_ was inversely related to the levels of ApoE3 but not correlated with ApoE4 levels (Edlund et al., 2021). Our findings are consistent with a report of very mild effects of *APOE* ε4 on GTT in human subjects (Virtanen et al., 2002).

We did not find an effect of ApoE variant on insulin resistance, consistent with results in humans (Meigs et al., 2000; Ragogna et al., 2012). Accumulating evidence suggests that metabolic disorders make a larger contribution to all-cause dementia in *APOE* ε4 noncarriers than in *APOE* ε4 carriers. For instance, *APOE* ε4 homozygotes with AD are much less likely to have hyperinsulinemia than AD patients with other genotypes (Craft et al., 1998). And ketogenic supplements, which provide a fuel source alternative to glucose, provide greater benefits to cerebral blood flow and cognition in *APOE* ε4 noncarriers(Henderson et al., 2009; Reger et al., 2004; Torosyan et al., 2018). The cognitive benefits of intranasal insulin are similarly exaggerated in *APOE* ε4 noncarriers (Reger et al., 2006). Thus, the cases of all-cause dementia among *APOE* ε4 noncarriers may include a greater number of vascular dementia, whereas *APOE* ε4 is associated with *bona fide* “pure” AD. Studies in which AD diagnosis is limited to neuropathologically confirmed cases have consistently found no association with type-2 diabetes or metabolic syndrome (Abner et al., 2016; Groeneveld et al., 2019; Hassing et al., 2002; Heyman et al., 1984; Peila et al., 2002; Sadrolashrafi et al., 2021; Thambisetty et al., 2013). Likewise, we find disparate effects of type-2 diabetes and AD on cerebral glucose transporters (manuscript in preparation).

One curious finding is that basal [Glc]_b_ seemed to be modulated in a manner distinct from the conditions which impaired glucose dynamics, i.e. GTT. The latter showed a significant effect only when Aβ expressers and controls were compared in the ApoE3 background. However, this scenario did not affect basal fasted [Glc]_b_, which was altered only by the combination of ApoE4 and Aβ accumulation. Multiple mechanisms of action have been attributed to ApoE, some of which involve lipid transport or other events dependent upon interaction of ApoE with its receptors. Distinct from these actions, ApoE4 can translocate to the cellular nucleus and act as a transcriptional repressor of a *cis* element termed coordinated lysosomal expression and regulation (CLEAR) (Lima et al., 2020; Parcon et al., 2018). This element is otherwise bound by the transcriptional inducer TFEB to elevate expression of several genes involved in autophagy and other lysosomal processes. Thus, *APOE* ε4 may elevate the accumulation of Aβ and the risk for AD by suppressing proteostasis generally (Raha et al., 2021). The maximal GTT values reached in Aβ-Tg mice were practically identical in mice expressing ApoE3 or ApoE4. This is reminiscent of the similar binding affinities that the two variants have for lipoprotein receptors (Weisgraber et al., 1982). (ApoE2, by contrast, exhibits a much weaker interaction with LDLR and related receptors.) It is tempting to speculate that the association of ApoE variant with glucose tolerance—essentially equivalent GTT results across the two variants—is a product of receptor binding, whereas the modality in which a substantial ApoE4-dependent effect was observed—ApoE4’s elevation of basal [Glc]_b_ in Aβ-expressing mice—is associated with a bioactivity in which a biochemical difference occurs, viz. CLEAR DNA binding. Failures in proteostasis appear to play an important role in the establishment of hyperglycemia and its underlying pathophysiology (Jung et al., 2008; Ozcan et al., 2009; Podolsky et al., 1972);Bachar-Wikstrom, 2013 #392}.

Human studies have consistently documented lower CMR_glc_ in subjects possessing one or more *APOE* ε4 gene, even at early ages (Reiman et al., 1996; Reiman et al., 1998; Small et al., 1995). An age-dependent reduction in glucose transport into the CNS has been reported in the ApoE-TR lines used here (Alata et al., 2015; Fang et al., 2021), though this was reported to occur at ages (8–12 months) far beyond most of the mice we examined. Surprisingly, another ApoE4-expressing model exhibits elevated CMR_glc_ as it progresses into advanced ages (Kotredes et al., 2021). ApoE4 is associated with structural and functional deficits in the cerebrovasculature (Tai et al., 2016; Zlokovic, 2013), and it is possible that this contributes to the impact of *APOE* genotype on CMR_glc_.

Effects of vascular conditions, including hyperglycemia, on CMR_glc_ typically involve downregulation of endothelial GLUT1 levels. Consistent with Alata et al. (2015)(Alata et al., 2015) and Wu et al. (2018)(Wu et al., 2018), we found no significant difference in total levels of GLUT1 (or other transporters) in ApoE4-TR mice. However, Aβ expression interfered with normal translocation of GLUT1 to the plasma membrane, as it does in mice with wild-type murine *Apoe* (Hendrix et al., 2020). This is correlated with reduced CMR_glc_ values, and we hypothesize that the insulin-independent glucose intolerance effected by Aβ represents a surrogate marker of the glucose which is not utilized in the CNS, due primarily to dysfunctional GLUT1. This hypothesis is supported by GTT results after conditional knockdown of GLUT1 in the astrocytes of mice (manuscript in preparation). It was surprising to see that cultured astrocytes were not affected in a manner predicted by *in vivo* results. Specifically, acute treatment of ApoE3-TR astrocytes with Aβ_1-42_ did not alter [^3^H]2DG as might be expected from the results with GTT and the PM fraction of 45-kDa GLUT1 *in vivo*. Instead, it was the ApoE4-TR astrocytes that exhibited diminished glucose transport in the presence of Aβ. Several differences between *in vivo* and *in vitro* conditions may explain this discrepancy, including the acute nature of the Aβ treatment *in vitro* and the cell-type complexity of the intact cerebrum.

IL-1β is well-documented as a component of neuroinflammation in AD. Previously, we found that application of inflammatory cytokines to cultured astrocytes could create the deficit in GLUT1 plasma-membrane trafficking seen in response to Aβ (Hendrix et al., 2020). Here, we found that IL-1β was elevated as a function of ApoE4 expression but not possession of the Aβ transgene. The mice used in this study were relatively young, particularly as regards plaque development in this model, which typically becomes detectable at ∼12 months of age.

In conclusion, we find that accumulation of Aβ_1-42_ impairs glucose transport and tolerance in mice expressing human ApoE3 but makes less, if any, impact on these parameters in the presence of ApoE4. Thus, it is possible that Aβ accumulation is less important for development of glucose perturbation in *APOE* ε4 carriers, and—to the extent that Tau pathology mirrored these relationships—it may be less important to dementing neuropathology. It is also notable that metabolic interventions such as diet-induced obesity have been reported to be exacerbated in ApoE4-TR mice, suggesting mechanistic differences in the glucose perturbations occurring in AD versus diabetes.

## Acknowledgments

This study was supported by funds from NIH/NIA (1R01AG071782, 1R01AG062254, and 2P01AG012411).

